# Two isoleucyl tRNAs that decode ‘synonymous’ codons divergently regulate breast cancer progression

**DOI:** 10.1101/2021.04.22.440519

**Authors:** Lisa B. Earnest-Noble, Dennis Hsu, Hosseinali Asgharian, Mandayam Nandan, Maria C. Passarelli, Hani Goodarzi, Sohail F. Tavazoie

## Abstract

The human genome contains 61 codons that encode for the 20 amino acids. The synonymous codons of a given amino acid are decoded by a set of transfer RNAs (tRNAs) called isoacceptors. We report the surprising observation that two isoacceptor tRNAs that decode synonymous codons are modulated in opposing directions during breast cancer progression. Specifically, tRNA^Ile^_UAU_ is upregulated, whereas tRNA^Ile^_GAU_ is repressed as breast cancer cells attained enhanced metastatic capacity. Functional studies revealed that tRNA^Ile^_UAU_ promoted and tRNA^Ile^_GAU_ suppressed metastatic colonization. The expression of these tRNAs mediated opposing effects on codon-dependent translation of growth promoting genes. Consistent with this, multiple mitotic gene sets in the human genome are significantly enriched in the codon cognate to the growth-promoting tRNA^Ile^_UAU_ and significantly depleted of the codon cognate to the growth-suppressive tRNA^Ile^_GAU_. Our findings uncover a specific isoacceptor tRNA pair that act in opposition—divergently regulating genes that contribute to growth and a disease phenotype. The degeneracy of the genetic code can thus be biologically exploited by human cancer cells via tRNA isoacceptor shifts that facilitate the transition towards a growth-promoting state.

## Main

Because of the degeneracy of the genetic code, multiple transfer RNAs (tRNAs) bearing distinct anticodons can accept the same amino acid for translational incorporation into the growing polypeptide chain during translation^1,2^. Such tRNA isoacceptors recognize what are called ‘synonymous codons’. Transfer RNAs have long been considered static adaptor molecules that play critical roles in converting the genetic code to an amino acid code. This notion has been revisited in recent years with observations of altered expression of tRNAs in the context of disease^3–5^, as well as demonstrated roles for certain over-expressed tRNAs (by genomic amplifications) as promoters of tumourigenic phenotypes^3,4,6^. Analogous to these observations, aminoacyl tRNA synthetases (aaRS), responsible for charging tRNAs with cognate amino acids, have been shown to play non-canonical roles^7^ and recent work has demonstrated significant cancer progression roles for specific charging enzymes^8,9^. These studies have raised a number of questions, including whether transcriptional deregulation in the absence of tRNA genomic copy number alterations can modulate tRNA levels and cancer progression as well as whether there exist metastasis suppressor tRNAs in human cancer.

### Isoleucyl tRNA isoacceptors are divergently modulated in breast cancer

To identify tRNAs that may become transcriptionally modulated during cancer progression, we performed chromatin immunoprecipitation sequencing (ChIP-seq) in poorly and highly metastatic human breast cancer cells^10^ using an antibody targeting the DNA binding subunit of Polymerase III, POLR3A. Enrichment of tRNA loci was confirmed by successful co-immunoprecipitation of Pol III genomic target loci through quantitative real-time PCR (qPCR) (Supplementary Fig. 1a), as well as significant enrichment of ChIP-seq reads for tRNA Box A and Box B gene regulatory sequences (Supplementary Fig. 1b). We observed that an isoleucyl-tRNA (TAT) isoacceptor locus that encodes tRNA^Ile^_UAU_ was significantly more bound by Pol III in highly metastatic MDA-LM2 cells relative to the parental poorly-metastatic MDA-MB-231 cells from which it was derived (Fig. 1a). To confirm these findings and to establish that mature tRNA^Ile^_UAU_ levels are upregulated in metastatic cells, we performed targeted tRNA profiling by tRNA Capture-seq^4^. Targeted tRNA quantification in the MDA-MB-231 poorly/highly metastatic pair as well as an independent poorly/highly metastatic isogenic human breast cancer line pair (HCC1806-Par and HCC1806-LM2C, validated in Supplementary Fig. 1c) confirmed that mature tRNA^Ile^_UAU_ is upregulated in highly metastatic breast cancer cells relative to isogenic poorly metastatic cells (Fig. 1b). Northern blot analysis confirmed the observations of tRNA^Ile^_UAU_ over-expression in highly metastatic breast cancer cells (Supplementary Fig. 1d). Genomic copy number analysis by qPCR did not reveal increased genomic copy number of isoleucyl-tRNA (TAT) loci in highly metastatic cells, consistent with transcriptional enhancement (Supplementary Fig. 1e). In parallel to these observations, we made the surprising observation that one of the other isoacceptors of isoleucine, tRNA^Ile^_GAU_, became significantly repressed in the highly metastatic sublines relative to isogenic poorly metastatic parental cells (Fig 1c). The high sequence similarity between tRNA^Ile^_GAU_ and another isoleucine isoacceptor tRNA^Ile^_AAU_ precluded specific northern blot quantification for tRNA^Ile^_GAU_ as an independent tRNA quantification method. We thus employed pre-tRNA quantification as an orthogonal approach for assessing the levels of all three isoleucyl tRNAs. Pre-tRNA qRT-PCR also revealed upregulation of tRNA^Ile^_UAU_ expression by the multiple genomic loci that encode it and conversely, repression of tRNA^Ile^_GAU_ loci genes in both pairs of highly metastatic breast cancer cells relative to their isogenic poorly metastatic parental cell populations (Fig 1d-e). We did not observe such global modulations of the third isoleucyl isoacceptor pre-tRNA^Ile^_AAU_ across the loci surveyed (Supplementary Fig. 1f). In support of these findings, FISH staining of human tissue microarrays of breast cancer patients with locked nucleic acids (LNAs) targeting tRNA^Ile^_UAU_ and tRNA^Ile^_GAU_ revealed a significantly increased ratio of tRNA^Ile^_UAU_/tRNA^Ile^_GAU_ expression in stage III breast tumours, which exhibit higher rates of metastatic relapse, relative to stage I or stage II tumours, which exhibit lower rates of metastasis (Fig. 1f). These findings reveal that metastatic progression in breast cancer selects for upregulation of one isoleucyl tRNA isoacceptor and repression of another. This shift in tRNA isoleucyl isoacceptor levels suggests potentially differential roles for these tRNAs in breast cancer progression.

**Figure 1.**
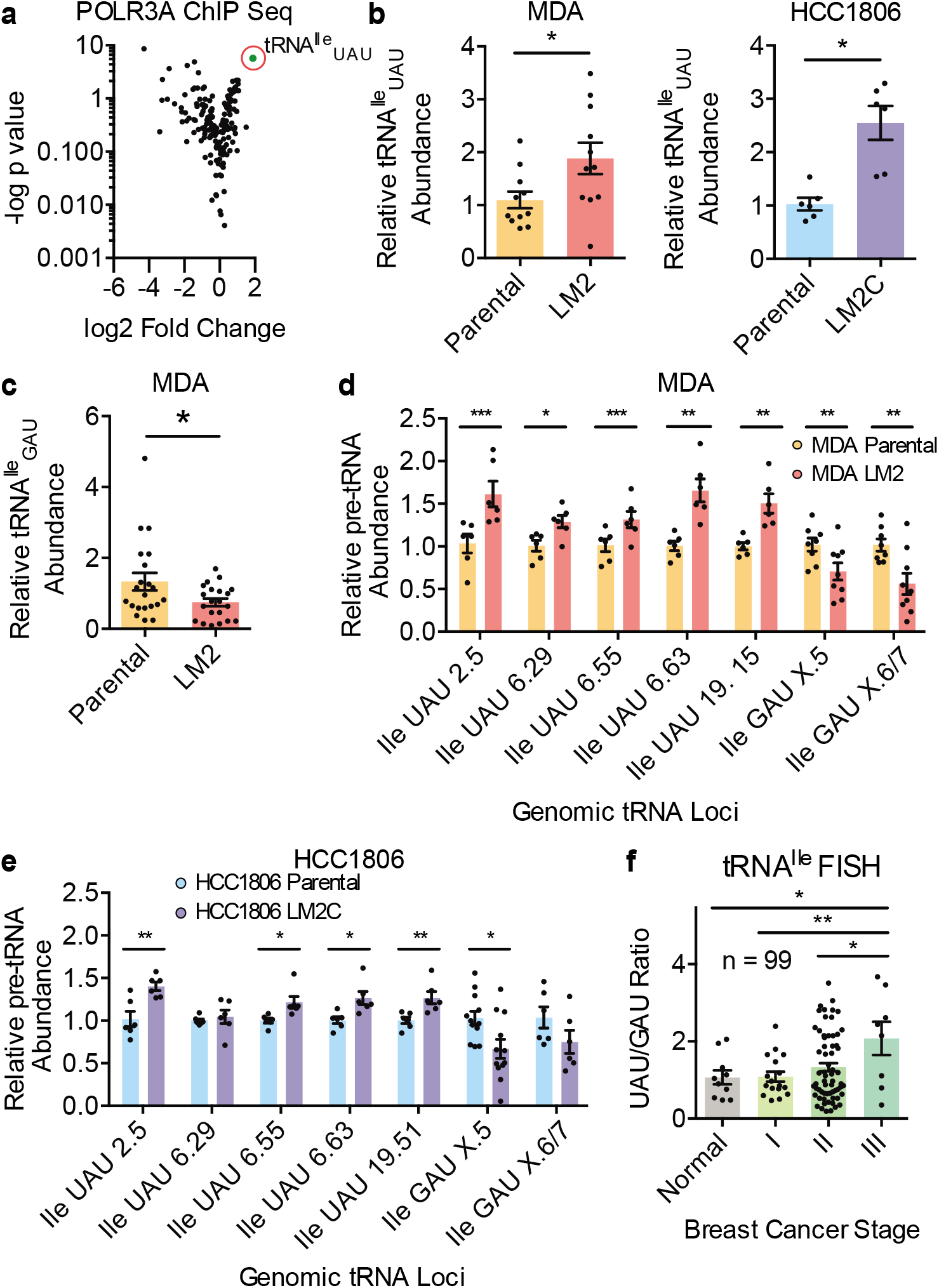
Isoleucine isoacceptors are differentially modulated in isogenic poorly and highly metastatic breast cancer pairs. (a) Volcano plot representing log2 fold change vs. –log p value of POL3RA ChIP sequencing analysis of MDA-LM2 cells vs. MDA-MB-231 Parental cells. (b) tRNA^Ile^_UAU_ quantification by specific tRNA^Ile^_UAU_ probe RT-qPCR normalized to 18S probes of highly metastatic LM2 lines relative to their parental MDA-MB-231 and HCC1806 cell lines. (c) tRNA^Ile^_GAU_ quantification by specific tRNA^Ile^_GAU_ probe RT-qPCR normalized to 18S probes of highly metastatic LM2 lines relative to the parental MDA-MB-231 cell line. (d,e) Relative pre-tRNA abundance of tRNA^Ile^_UAU_ and tRNA^Ile^_GAU_ across multiple primers covering distinct genetic loci using RT-qPCR of MDA-LM2 vs. MDA-MB-231 (d) & HCC1806-LM2C vs. HCC1806 Parental cells (e). (f) Relative tRNA^Ile^_UAU_/tRNA^Ile^_GAU_ ratios quantified by fluorescent intensity normalized to DAPI of breast tissue microarrays, stratified by normal tissue or breast cancer stage I & II, III, measured by FISH with LNA targeting tRNA^Ile^_UAU_ or tRNA^Ile^_GAU_. Two-sided un-paired student’s t-tests performed, p-values p<0.05, p<0.01, p<0.001 represented as *, **, ***, respectively.

### TRNA^Ile^_UAU_ promotes and tRNA^Ile^_GAU_ suppresses breast cancer metastasis

To determine if the observed reciprocal tRNA isoleucyl isoacceptor modulations play causal roles in cancer progression, we performed loss-of-function and gain-of-function studies for these tRNA isoacceptors. We first sought to overexpress tRNA^Ile^_UAU_ in poorly metastatic cells to assess whether its upregulation was sufficient to confer increased metastatic capacity (Supplementary Fig. 2a-b). Stable over-expression of tRNA^Ile^_UAU_ to pathophysiologically relevant levels (~50% increase) in poorly metastatic MDA-MB-231 or HCC1806 human cell lines significantly increased lung metastatic colonization in tail-vein colonization assays as assessed by bioluminescence imaging and histological analyses (Fig. 2a-b). For loss-of-function studies, we employed CRISPR-Cas9 using two independent guides specific to tRNA^Ile^_UAU_ genomic loci (Supplementary Fig. 2c). CRISPR-Cas9 mediated depletion of tRNA^Ile^_UAU_ in highly metastatic MDA-LM2 breast cancer cells to levels similar to poorly metastatic cells was sufficient to significantly impair breast cancer metastatic colonization (Fig. 2c). These findings reveal tRNA^Ile^_UAU_ to be a promoter of metastatic progression in these human breast cancer cells.

**Figure 2.**
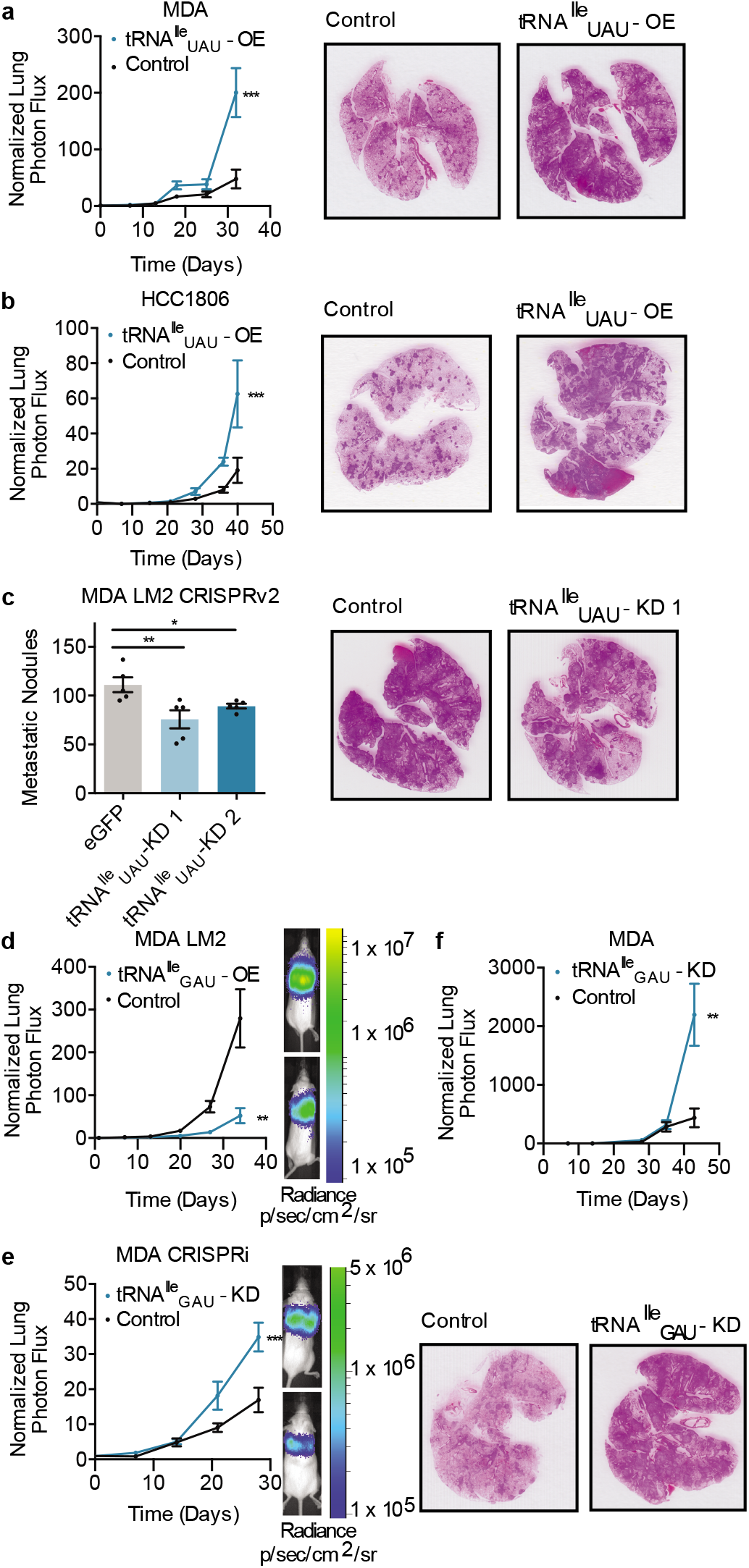
tRNA^Ile^_UAU_ promotes & tRNA^Ile^_GAU_ suppresses metastatic colonization. (a-b) Bioluminescent imaging post tail vein injection of 1×10^5^ of MDA Parental (a) or 1.5×10^5^ HCC1806 (b) cells overexpressing tRNA^Ile^_UAU_ or control with representative lung histology stained with H&E; n=5 in each cohort. (c) Quantification of lung metastatic nodules post extraction after tail vein injection of 5×10^4^ LM2 CRISPR cells guides targeting eGFP or tRNA^Ile^_UAU_, with representative histology for control & tRNA^Ile^_UAU_ guide 1; n=5 in each cohort. (d) Bioluminescence imaging after tail vein injection of 1×10^5^ of MDA LM2 cells overexpressing tRNA^Ile^_GAU_ or control with representative images; luminescence expressed as Radiance p/sec/cm^2^/sr; n=5 in each cohort. (e,f) Same as (d) with MDA Parental CRISPRi cells with guides targeting either control or tRNA^Ile^_GAU_ (e) or shRNA targeting control or tRNA^Ile^_GAU_ with representing H&E lung histology (f). Statistics utilized include 2-way ANOVA for imaging and two-sided unpaired student’s t-test for nodule quantification, p-values p<0.05, p<0.01, p<0.001 represented as *, **, ***, respectively.

We next determined if tRNA^Ile^_GAU_, which became repressed in metastatic cells, plays a causal role in breast cancer progression. TRNA^Ile^_GAU_ was stably overexpressed in highly metastatic MDA-LM2 cells to levels similar to those observed in poorly metastatic MDA-231 parental cells (~1.8-fold over-expression) (Supplementary Fig. 2d). Increasing tRNA^Ile^_GAU_ expression in highly metastatic MDA-LM2 cells substantially reduced metastatic lung colonization capacity (Fig. 2d). Given the high sequence similarity between tRNA^Ile^_GAU_ and tRNA^Ile^_AAU_, we employed two orthogonal approaches for tRNA^Ile^_GAU_ loss-of-function—CRISPRi and shRNA mediated interference. Firstly, MDA-231 cells were stably transduced with mutant Cas9-KRAB and a specific guide complementary to common sequences in tRNA^Ile^_GAU_ genomic loci. Reduced tRNA^Ile^_GAU_ was confirmed by targeted tRNA capture qPCR (Supplementary Fig. 2e). ShRNA-mediated interference was also employed using a hairpin specific to tRNA^ile^_GAU_ (Supplementary Fig. 2f). Depletion of tRNA^Ile^_GAU_ using both approaches enhanced lung metastatic colonization by poorly metastatic MDA-231 cells (Fig. 2e and 2f). These findings implicate tRNA^Ile^_GAU_ as a metastasis suppressor tRNA and uncover two surprising findings: the first being a gain-of-function organismal disease phenotype upon depletion of a tRNA (tRNA^Ile^_GAU_); the second being the observation of a dichotomy between two tRNA isoacceptors in regulating a common phenotype.

### TRNA^Ile^ isoacceptors divergently regulate growth and growth gene expression

We next sought to identify the cancer progression cellular phenotype(s) regulated by tRNA^Ile^_UAU_ and tRNA^Ile^_GAU_ by searching for gene sets that exhibit enrichments or depletions of codons cognate to these tRNAs. We performed pathway enrichment analyses using the iPAGE framework^11^—assessing significant genome-wide abundances for the codons cognate to tRNA^Ile^_UAU_ and tRNA^Ile^_GAU_ in an unbiased manner. All coding transcripts in the human genome were ranked and binned by AUA or AUC relative synonymous codon usage (RSCU). Pathways that were significantly enriched (p<10^-3^) across discretized bins were identified based on their mutual information content (Fig. 3a-b). Interestingly, transcripts most signficantly enriched in AUA codons (cognate to tRNA^Ile^_UAU_) were enriched in mitosis related gene sets such as metaphase, anaphase, and chromatid separation, and homologous DNA pairing and strand exchange (Fig. 3a). Conversely, AUC codons (cognate to tRNA^Ile^_GAU_) were most significantly depleted from these mitosis related gene sets (Fig. 3b). As an orthogonal and functional approach for identifying the downstream consequences of modulation of these tRNAs, we conducted ribosomal profiling of breast cancer cells in the context of tRNA^Ile^_UAU_ overexpression and tRNA^Ile^_GAU_ depletion (by CRISPRi), mirroring the divergent tRNA^Ile^ modulations observed in highly metastatic cells relative to poorly metastatic cells. Ribosomal protected fragments were sequenced, and conformed to the expected size and periodicity reported by other groups^12^ (Supplementary Fig. 3a-b). Ribosomal occupancy of transcripts was then quantified as a measure of translational efficiency (Supplementary Fig 3c). Genes enriched in GO terms such as cell cycle and mitosis exhibited enhanced translational efficiency (Fig. 3c). At the proteomic level, GO functional analysis of proteins in tRNA^Ile^_UAU_/tRNA^Ile^_GAU_ modulated cells by label free mass spectrometric quantification also revealed enrichment of gene sets including cell cycle, mitosis, as well as regulation of stress response relative to control cells (Fig. 3d). Consistent with the growth related gene sets identified using the described approaches, immunofluorescent staining of metastatic nodules for the proliferation marker Ki67 revealed that MDA MB 231 breast cancer cells concomitantly over-expressing tRNA^Ile^_UAU_ and depleted of tRNA^Ile^_GAU_ exhibited greater proliferation than control cells (Fig. 3e, Supplementary Fig. 3d). To determine if these *in vivo* observations could be recapitulated *in vitro*, growth assays were performed under normal tissue culture conditions and under conditions of hypoxic and oxidative stress, since such stresses occur in the metastatic microenvironment and can restrict growth^13–18^. Concomitant tRNA^Ile^_UAU_ upregulation/tRNA^Ile^_GAU_ depletion enhanced the *in vitro* growth of MDA MB 231 breast cancer cells relative to control cells in the context of hypoxia (Fig. 3f) and oxidative stress (Fig. 3g). Importantly, growth effects were more pronounced under these stress conditions that are known to occur in the tumour microenvironment than under normoxic basal *in vitro* conditions (Supplementary Fig. 3e). These findings reveal that divergent modulation of these isoleucyl tRNA isoacceptors promotes growth in these breast cancer cells *in vivo* and *in vitro*.

**Figure 3.**
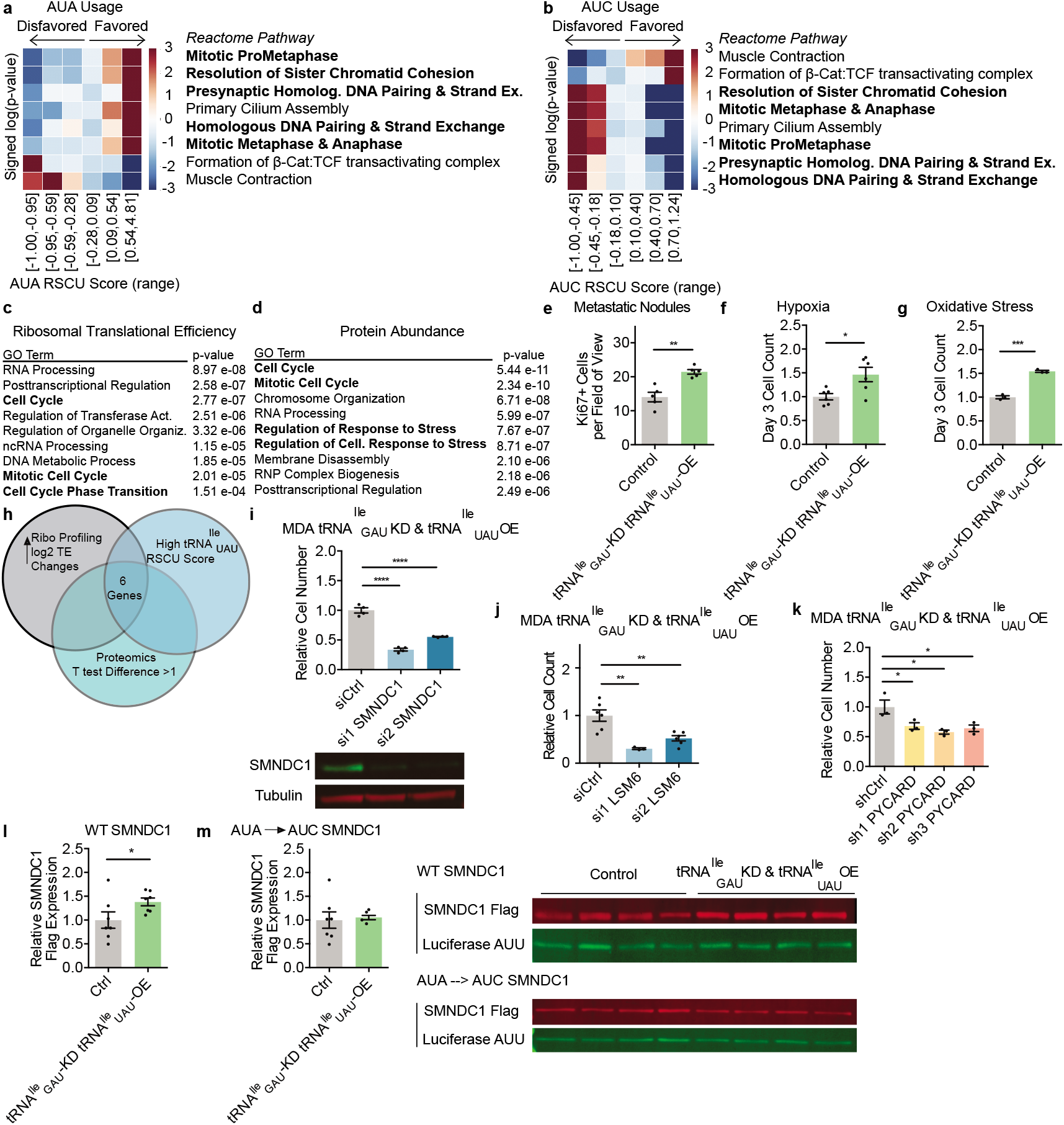
Cell Cycle and Response to Stress gene expression and phenotypes characterize tRNA^Ile^ modulations. (a,b) Reactome pathways significantly enriched in AUA (a) or AUC (b) by relative synonymous codon usage (RSCU) using iPAGE. (c,d) GO function terms for positive and significant TE changes from ribosomal profiling (c) or label free quantification by mass spectrometry (d) in tRNA^Ile^_GAU_ depletion and tRNA^Ile^_UAU_ overexpression cells versus control. (e) Quantification of Ki67 immunofluorescence staining in MDA MB 231 tRNA^Ile^_GAU_ depletion and tRNA^Ile^_UAU_ overexpression cells versus control. (f,g) Relative cell counts of MDA MB 231 control & tRNA^Ile^_GAU_ depletion tRNA^Ile^_UAU_ overexpression cells exposed to 0.5% hypoxia (f) or treated with 200uM H202 (g) for 3 days. (h) Venn Diagram of overlapping datasets to identify downstream effectors – includes high RSCU tRNA^Ile^_UAU_ score (top 50%), and genes with significantly positive changes in TE and proteomics in both MDA LM2 vs. MDA Parental cells and tRNA^Ile^_GAU_ depletion tRNA^Ile^_UAU_ overexpression cells vs. control. (i) Relative cell count of MDA tRNA^Ile^_GAU_ depletion tRNA^Ile^_UAU_ overexpression cells treated with control or SMNDC1 siRNA in 0.5% hypoxia for 2 days (d). Western performed on siRNA cells on day 3. (j) Relative cell counts of MDA tRNA^Ile^_GAU_ depletion tRNA^Ile^_UAU_ overexpression cells treated with control or LSM6 siRNA for 3 days. (k) Relative cell counts of MDA tRNA^Ile^_GAU_ depletion tRNA^Ile^_UAU_ overexpression cells transduced with shRNA targeting either control or PYCARD for 3 days. (l,m) LICOR Western quantification of either Flag tagged wildtype (l) or all AUA to AUC codons (m) SMNDC1 expression relative to reporter control luciferase (all Ile AUU) 24 hours post transfection in either MDA control or tRNA^Ile^_GAU_ depletion tRNA^Ile^_UAU_ overexpression cells. Representative images below. Statistics utilized include two-sided un-paired student’s’s t-tests performed, p-values *, **, **** indicated as p<0.05, p<0.01, and p<0.0001, respectively.

### A growth gene network regulated by TRNA^Ile^ isoacceptors

We next sought to identify examples of downstream effector genes that could mediate cell growth effects downstream of tRNA^Ile^_UAU_/tRNA^Ile^_GAU_ modulation. We hypothesized that there exist growth-promoting genes enriched in AUA codons cognate to tRNA^Ile^_UAU_. We identified the set of genes that exhibited enhanced translational efficiency as well as enhanced mass-spectrometric protein abundances upon concurrent tRNA^Ile^_UAU_/tRNA^Ile^_GAU_ modulation, and exhibited a high relative synonymous codon usage score for tRNA^Ile^_UAU_. The ten genes that fulfilled these criteria were further restricted to those that exhibited enhanced translational efficiencies and protein abundances in highly metastatic cells, which endogenously modulate these tRNAs relative to the isogenic parental poorly metastatic population (Fig. 3h)^4^. This yielded six genes as candidate downstream growth-promoting effectors (Supplementary Fig. 3f). Functional testing revealed that RNAi-mediated depletion of three of these genes (*SMNDC1, LSM6*, and *PYCARD*) reduced proliferation (3i-k, Supplementary Fig. 3g, h). We next focused on one gene, *SMNDC1*, for mutagenesis studies. To determine if isoleucyl tRNA modulations can directly enhance translation of a growth-promoting gene in a codon-dependent manner, we employed a reporter-based approach in which AUA codons in *SMNDC1* were mutated to synonymous AUC codons. While the wildtype SMNDC1 protein became upregulated upon dual tRNA modulation, synonymous codon mutant SMNDC1 protein levels remained unchanged (Fig. 3l, m)—consistent with codon-dependent tRNA^Ile^_UAU_-driven enhancement of translation of this growth-promoting gene. These findings reveal that divergent isoleucyl tRNA modulation enhances translation of a set of growth-promoting genes with high tRNA^Ile^_UAU_ relative synonymous codon usage scores.

### Divergent tRNA isoacceptor modulation impacts ribosomal function

The opposing directionality of the metastasis phenotype observed upon modulating these isoacceptor tRNAs suggests that they may elicit distinct downstream codon-dependent translational effects at a global level. To test this, we performed polysome profiling studies (Supplementary Fig. 4a). This revealed that relative to control cells, concurrent tRNA^Ile^_UAU_ over-expression and tRNA^Ile^_GAU_ depletion elicited a significant increase in polysome occupancy of transcripts enriched in the AUA codon, which is cognate to the over-expressed tRNA^Ile^_UAU_ (z-score 23.6; robustness 10/10; Fig. 4a) and a reduction in actively translating transcripts enriched in the AUC codon, which is cognate to the depleted tRNA^Ile^_GAU_ (z-score 44.8; robustness 10/10; Fig. 4b). Consistent with this, analysis of the aforementioned ribosomal profiling data revealed that upon dual tRNA^Ile^_UAU_/tRNA^ile^_GAU_ modulation, there was also a significant enrichment of ribosomal occupancy of AUA-containing transcripts and reduced occupancy of AUC-containing transcripts (Supplementary Fig. 4b,c). Our findings as a whole suggest a model whereby tRNA^Ile^_UAU_/tRNA^Ile^_GAU_ modulation enhances the efficiency of AUA codon decoding by the ribosome. This would suggest that we should observe reduced ribosomal dwell time over AUA codons upon dual tRNA modulation. Moreover, we would expect to see increased binding of tRNA^Ile^_UAU_ relative to tRNA^Ile^_GAU_ to the ribosome upon tRNA^Ile^_UAU_/tRNA^Ile^_GAU_ modulation. In order to capture the dwell time of ribosome at every codon, we measured the extent to which its occupancy in the ribosome profiling data deviates from its predicted level based on loess regression^19^. We observed that tRNA^Ile^_UAU_/tRNA^Ile^_GAU_ modulation significantly reduced ribosome dwell time over AUA codons, consistent with productive translation, while over-expression of tRNA^Ile^_UAU_ or depletion of tRNA^Ile^_GAU_ individually were insufficient to elicit significant shifts in dwell time (Fig. 4c). To determine if tRNA modulations impact ribosome-associated tRNA^Ile^_UAU_ and tRNA^Ile^_GAU_ abundances, we quantified the abundance of these tRNAs from polysomal ribosomes as well as total cellular input. We observed that tRNA^Ile^_GAU_ depletion reduced the ribosomal association of this tRNA, while tRNA^Ile^_UAU_ over-expression enhanced its ribosome association (Fig. 4f). Importantly, dual tRNA^Ile^_UAU_/tRNA^ile^_GAU_ modulation caused the greatest increase in relative tRNA^Ile^_UAU_ to tRNA^Ile^_GAU_ ribosomal association (Fig. 4g). The substantially increased ribosomal association of tRNA^Ile^_UAU_ upon dual tRNA^Ile^_UAU_/tRNA^Ile^_GAU_ modulation relative to tRNA^Ile^_GAU_ depletion supports the translational consequences observed upon polysome profiling (Fig. 4d, e). These concordant observations of global shifts in isoleucine codon enrichments and depletions in polysome profiling and ribosomal profiling studies as well as dwell time and biochemical analyses support direct codon-dependent effects on translation upon divergent modulation of these tRNAs. Our findings as a whole support a model whereby isoleucyl isoacceptor tRNA abundance shifts impact codon-dependent translation of growth regulating genes at the ribosome, thereby promoting cancer progression (Fig. 4h).

**Figure 4.**
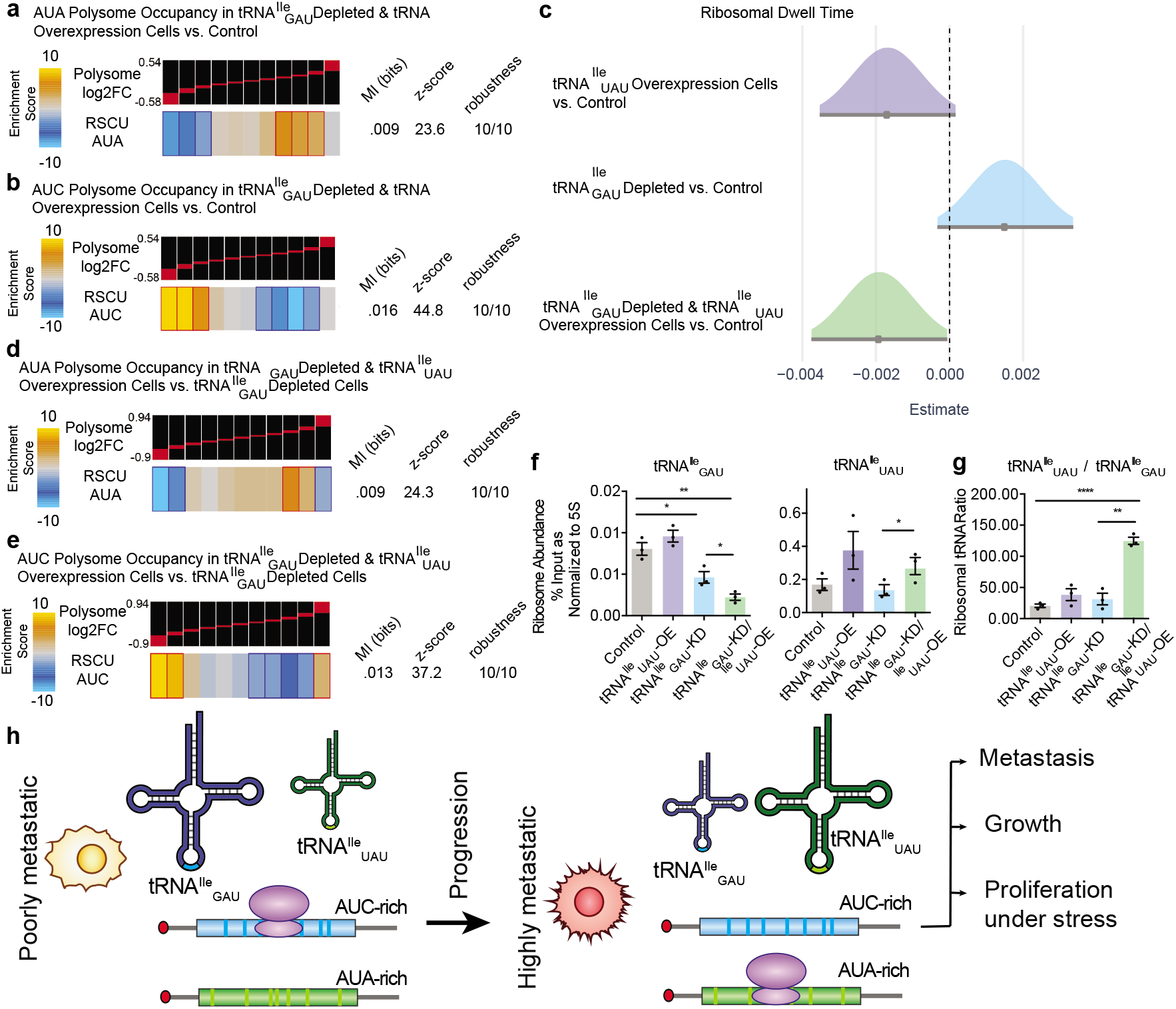
Translational efficiency of AUA enriched transcripts is dependent on tRNA^Ile^_GAU_ abundance. (a) Genes with a high abundance of AUA codons (using RSCU scores) were significantly enriched among genes upregulated in polysomes (corrected for their transcript changes) in tRNA^Ile^_GAU_ depleted tRNA^Ile^_UAU_ overexpression cells versus control MDA-MB-231 cells. The statistical significance of these enrichments was assessed using mutual-information calculations and associated Z score (based on randomized input vectors) and robustness scores (based on jackknifing tests). The heatmap was generated using the –log of the hypergeometric p-value for enrichment and log of p-value for depletion (collectively termed the enrichment score). The red and dark-blue borders indicate the statistical significance of the calculated hypergeometric p-values (for details, see Goodarzi et al., 2009)^11^. (b) Same as (a) except analyzed for AUC codon enrichment, showing significantly depletion among genes upregulated in polysomes (corrected for their transcript changes) in tRNA^Ile^_GAU_ depleted tRNA^Ile^_UAU_ overexpression cells versus control MDA-MB-231 cells. (c) Ribosomal AUA codon dwelling times as estimated by CELP bias coefficients (higher bias coefficient indicates longer dwelling time). Univariate regression coefficients estimating the effects of tRNA^Ile^ modulated MDA cells. A 95% confidence interval excluding zero (not overlapping the vertical line x=0) means that the tested effect was significant at α=0.05 (p<0.05). (d) Hypergeometric distribution shown as a z-score of AUA codon enrichment of polysome transcripts represented as log2 fold change of tRNA^Ile^_GAU_ depleted tRNA^Ile^_UAU_ overexpression cells versus tRNA^Ile^_GAU_ depleted MDA MB 231 cells, stratified in bins of 10, increased log2 fold change from left to right. AUA codon representation visualized as a value ranging from −10 to 10 relative to the average. (e) Same as (d) except analyzed for AUC codon enrichment. (f) Ribosome abundance of tRNA quantified by specific tRNA GAU (left) and tRNA UAU (right) probes. RT-qPCR normalized to 5S probes of tRNA modulated MDA cells from polysome fractions, measured as % Input. (g) Ribosomal ratio of tRNA^Ile^_UAU_/tRNA^Ile^_GAU_ abundance quantified by specific tRNA^Ile^_GAU_ probe and tRNA^Ile^_UAU_ probe RT-qPCR normalized to 5S probes of tRNA^Ile^ modulated MDA cells from polysome fractions, measured as % Input. (h) Model depicting how tRNA^Ile^ abundance shifts alter translational dynamics and metastatic phenotypes. Two-sided un-paired student’s’s t-tests performed, p-values represented as *, ** as p<0.05, p<0.01 respectively.

## Discussion

Our observations reveal opposing roles for two isoleucyl tRNAs in regulation of breast cancer metastatic colonization and cancer cell growth. Our findings as a whole support a model whereby shifts in tRNA^Ile^ isoacceptor abundance impact codon-dependent translation of growth regulating genes at the ribosome, thereby promoting cancer progression (Fig. 4h). The molecular and functional studies implicating growth as a phenotype divergently impacted by modulation of these tRNAs is supported by genome sequence analyses that reveal significant enrichment or depletion of the codons cognate to these antagonistic tRNAs in mitotic gene sets. Future studies are warranted to better elucidate the molecular basis of such interferences and to search for additional examples of such antagonistic isoacceptor tRNA pairs in health and disease.

## Methods

### Cell Culture

MDA-MB-231 and its highly metastatic derivative^10^ LM2 cells were cultured with DMEM media supplemented with 10% FBS, sodium pyruvate, and L-glutamine. HCC1806 Parental and derivate cell lines were cultured in 1x RPMI supplemented with 10% FBS, sodium pyruvate, 1mM HEPES as specified by ATCC. All cell lines were regularly tested for mycoplasma infection and were negative. Each cell line was verified using STR testing, performed by the Integrated Genomics Operation at MSKCC. Cells were retrovirally transduced with a luciferase reporter for bioluminescence detection as previously described^4,15,20^. Oxidative stress analyses were conducted by addition of 200 uM hydrogen peroxide to cells.

### *In Vivo* Selection

Several female Nod SCID Gamma (NSG) (Jackson # 005557) mice were injected at 6 weeks of age intravenously via tail vein with 150,000 parental HCC1806 cells and monitored by bioluminescence IVIS imaging (IVIS Lumina II) until photon flux of lungs reached 10^7 or 10^8 (4-7 weeks). Subsequently, animals were euthanized according to IACUC protocol and guidelines, and the lungs were extracted under sterile conditions. The lungs were then placed on a sterile 6cm tissue culture dish and minced with razor blades. Lung tissue extracts were re-suspended in 20 mL RPMI Media supplemented with FBS, sodium pyruvate and HEPES with 15 mg/mL Collagenase IV (Worthington). Cells were incubated at 37°C for 30 minutes on a shaker to allow for digestion. Tissue extracts were then spun down at 1000 RPM for 5 minutes at 4°C and resuspended in RPMI media without Collagenase IV. Cells were filtered with a 100 um filter (Corning) and spun down again. Cells were then treated with 5 mL ACK Lysis Buffer and left at room temperature for 5 minutes. Cells were spun again and resuspended in 1 mL Optiprep solution 1 (2:1, Optiprep:Media). A gradient was constructed with an additional 4 mL Solution 1, overlain by 3 mL Optiprep Solution 2 (2.2:1, Solution1: Media). 1 mL Media was overlain and the gradient was spun down for 20 minutes at 1000 RPM 4°C. Viable cells were collected from the top of the gradient and washed twice with RPMI media. Cells were then plated in 75 cm filtered flasks with PenStrep and Fungizone added to the RPMI Media. The cells were cultured for approximately a week to reduce stromal cell survival and then tested for mycoplasma. To generate an in vivo selected line twice, these lung metastatic 1 (LM1) generation cells were then re-injected at 150,000 tail vein and the process was repeated.

### Chromatin Immunoprecipitation

MDA-MB-231 and LM2 cells in biological replicates were plated in 15 cm plates (~12 million cells). For cross-linking, 1% formaldehyde was added to cells at 37°C for 10 minutes, and quenched with glycine at a final concentration of 0.14 M for 30 minutes at room temperature. The plates were put on ice, the media was removed and cells were washed with ice cold PBS twice. 500 uL PBS with 1x HALT protease inhibitors (Thermo) were added and cells were scraped and put in an Eppendorf on ice. Cells were pelleted at 4000 RPM in a refrigerated centrifuge for 4 minutes. The cell pellets were then resuspended in 400 uL Lysis buffer (1% SDS, 50 mM Tris Hcl pH 8.0 20 mM EDTA, protease inhibitors (Roche) and incubated on ice for 10 minutes. Lysates were then sonicated to produce DNA fragments between 200 – 1000 bp with settings Amplitude 70, 10 sec on 30 sec rest for three repetitions on Sonicator S-4000 (Branson) with Microtip and Ultrasonic Liquid Processor (Misonix). Lysates were kept on ice during sonication to prevent protein degradation. Lysates were then clarified by centrifugation at max speed at 4 °C for 10 minutes. Equivalent amounts of lysate were then added to separate eppendorfs, saving some lysate for input samples. Either 5 ug rabbit IgG or 5ug POLR3A (Cell Signaling #12825S) were then added, with a final volume of 1 ml with protease inhibitors of dilution buffer (16.7 mM Tris HCl pH 8.0, 0.01% SDS, 1.1% Triton X-100, 1.2 mM EDTA, 167 mM NaCl). Lysate and antibody mixtures were incubated overnight at 4 °C with rotation. 50 uL Protein G Dynabeads were added to each sample after washing and incubated at 4 °C for 2 hours with rotation. Tubes were then placed on a magnet for 2 minutes, discarding the supernatant. The following washes were performed, twice each for 5 minutes at 4 °C in the following order: low salt (140 mM NaCl, 50 mM HEPES, 0.1% SDS, 1% Triton X-100, 0.1% deoxycholate, 1 mM EDTA) high salt (500 mM NaCl, 50 mM HEPES, 0.1% SDS, 1% Triton X-100, 0.1% deoxycholate, 1 mM EDTA), LiCl (250 mM LiCl, 20 mM Tris HCl pH 8.0, 0.5% NP-40, 0.5% deoxycholate, 1 mM EDTA) and TE Buffer (10 mM Tris HCl pH 8.0, 1 mM EDTA). 100 uL Elution buffer (50 mM Tris HCl pH 8.0, 1 mM EDTA) was then added to the beads and incubated overnight at 65 °C on a shaker to enable elution. The eluted sample was transferred to a new tube and repeated for a final volume of 200 uL per sample. 1 uL of 10 mg/mL RNase A was added to each sample (including input samples) and incubated at 37 °C for 30 minutes. 2 uL of Proteinase K was added (10 mg/mL) and incubated for 2 hours at 56 °C. DNA was then purified using the DNA Clean and Concentrator Kit (Zymo Research). Enrichment of Polymerase III bound loci was confirmed with genomic tRNA qPCR primers and quantified as Percent Input over IgG. The Input and IP DNA samples were then PCR amplified with Illumina barcodes to construct a multiplexed library. The library was quantified using TapeStation and sequenced on the 50 SR HiSeq at the Rockefeller Genomics Resource Center.

### tRNA Capture qPCR

tRNA quantification by RT-qPCR was performed as described previously^1^. RNA purified with Norgen Total Purification Kit was quantified using a Nanodrop. Normalized RNA across samples was added to a hybridization mixture (final concentration 10 mM Tris HCl pH 7.4, 50 mM NaCl, 1 mM EGTA pH 8.0) with 2 uM Hybridization probes. Hybridization probes specific for the following tRNA were used: Ile TAT Left 5’/5Phos/AAGTACCGCGCGCTACCGATTGCGCCACTGGAGCGATCGTCGGACTGTAGAA, Ile TAT Right 5 CGTGTGCTCTTCCGATCTTGCTCCAGGTGAGGCTCGAACTCACACCTCGGCATTAT’, Ile GAT Left 5’/5Phos/CAGCACCACGCTCTACCAACTGAGCTAACCGGCCGATCGTCGGACTGTAGAA, Ile GAT R 5’ CGTGTGCTCTTCCGATCTTGGCCGGTGCGGGAGTCGAGCCCGCGCCTTGG TGTTAT 3’. Each ‘left’ probe contained a 5’ phosphate to enable subsequent ligation. RNA and probe mixture was hybridized using a thermocycler and brought to RT. 1x SplintR ligase buffer, SplintR Ligase (NEB) and RNase Inhibitor (Promega) were added and incubated at room temperature for 2 hours. An additional ligation step with T4 Ligase was performed overnight at 16 °C. The RNA was then degraded with RNase A (Thermo Fisher) & H (NEB) for 30 minutes at 37 °C. The ligated probe reaction was then diluted 1:50 and quantified using primers (Forward 5’ CGTGTGCTCTTCCGATCT 3’ & Reverse 5’ GATCGTCGGACTGTAGAA 3’) specific to the probe backbone by RT-qPCR. 5S and 18S probes were used as loading controls: 5S Left 5’ 5PHOS/CTGCTTAGCTTCCGAGATCAGACGAGATCGGGCGCGATCGTCGGACTGTAGAA 3’ 5S Right 5’ CGTGTGCTCTTCCGATCTCCAGGCGGTCTCCCATCCAAGTACTAACCAGGCCCGACC 3’ or 18S Left 5’ 5PHOS/CCTAGTAGCGACGGGCGGTGTGTACAAAGGGCGCCGATCGTCGGA CTGTAG 3’ 18S Right 5’ CGTGTGCTCTTCCGATCTCCGATCCGAGGGCCTCACTAAACCATCCAATC 3’.

### Northern Blot

RNA was purified using Norgen total RNA Purification kits according to manufacturer’s instructions. 5 ug purified RNA was run on 10% TBE-Urea gels at 200V for 1 hour, and transferred to a Hybond-N+ membrane (GE) at 150A for 1 hour. RNA was then crosslinked to the membrane at 240 mJ/cm^2^, and blocked with Oligo Hybridization Buffer (Ambion) for 1 hour at 42 °C. Northern probes were labeled with ^32^P ATP with T4 PNK (NEB), purified with a G25 column (GE Healthcare), and hybridized in Oligo Hybridization Buffer overnight at 42 °C. Membranes were washed with 2X SSC 0.1% SDS Buffer, then with 1X SSC 0.1% SDS Buffer. Films were developed at varying times subject to radioactivity of membrane. Probe oligo sequences for Ile TAT Intron: 5’ ACUGCUGUAUAAGUACCGCGCGC 3’ and Ile TAT 5’ cucggcauuauaaguaccgcgcgc 3’ and U6 5’ CACGAATTTGCGTGTCATCCTT 3’. Membranes were stripped with 0.1% SDS in boiling water and allowed to cool to room temperature. Quantification was performed with ImageJ and normalized to U6 levels.

### RT qPCR

RNA was purified using the Norgen total RNA Purification kits according to manufacturer’s instructions. 1ug purified RNA was used for cDNA production with Superscript III reverse transcriptase (Thermo Fisher Scientific) using random hexamer as a template. The cDNA was diluted 1:5 and quantified with Sybr Green Master Mix (Thermo). The ddCT levels were quantified through normalization to 18S with biological replicates. One primer set was used for tRNA^Ile^_GAT_ genetic loci chr.X-6 and chr.X-7 as their sequences are indistinguishable. Primer sequences are available in the Supplemental Material Section.

### tRNA Fluorescence In Situ Hybridization

Breast tissue microarrays were obtained from Biomax (BC08118 & BR1005b). Slides underwent deparaffination via 5 minute incubations in xylene 2x, 100% Ethanol, 2x, 70% Ethanol, 50% Ethanol, and subsequently molecular grade water. Antigen retrieval was performed with 1x Citrate Buffer pH 6.0 in a microwave for 20 minutes. Slides were cooled to room temperature, then tissue regions were isolated with a PAP pen. Slides were incubated in 0.13 M 1-methylimidizaole 300 mM NaCl pH 8.0 solution twice for 10 minutes each. Next, slides were incubated with 0.16 M N-(3-Dimethylaminopropyl)-N-ethylcarbodiimide hydrochloride (EDC) (Sigma Aldrich) in the 1-methylimidazole solution for 1 hour at RT to preserve small RNAs^21^. Slides were then washed with 0.2% Glycine in Tris buffered saline (TBS) pH 7.4, then in TBS twice. Pre-hybridization of slides occurred with 1X ISH (Exiqon) buffer at 53 °C for 1 hour. 40 nM tRNA double DIG labeled LNA Probe targeting tRNA^Ile^_UAU_ (Sequence 5’ CA+GGTGAGGCTCGAACTCACAC+C+TCGGCAT+T+A 3’ with +N indicating LNA at that nucleotide) and tRNA^Ile^_GAU_ (Sequence 5’ AGTCGA+GCCCGCGAC+CTTGG+TGTTA+T+C 3’) (Qiagen) in 1X ISH buffer was denatured at 95°C for 5 minutes followed by cooling on ice for 1 minute. The LNA probe was added to the slide (and covered with a glass coverslip to prevent evaporation) and hybridized overnight at 52 °C. Slides were then washed with 4X SSC, 2X SSC, 1X SSC, and 0.5X SSC in 50% formamide for 20 minutes each, then washed with 100 mM Tris-HCl pH 7.4 150 mM NaCl (TN buffer) for 5 minutes. Slides were blocked with 1X Blocking Reagent (Roche) in TN buffer for 1 hour at RT. Anti-DIG POD in TN blocking buffer was added 1:100 and incubated for 2 hrs. Slides were washed 3x for 5 minutes in TN buffer with 0.05% Tween-20 (TNT). FITC-tyramide solution 1:100 in 1x amplification reagent (TSA) was incubated on slides for 10 minutes at RT. Slides were subsequently washed 3x for 5min with TNT buffer. Samples were washed with PBS and stained with DAPI for 5 minutes, then mounted with Prolong Gold anti-fade solution (Thermo Fisher). Fluorescent intensity was measured on an Inverted TCS SP8 laser scanning confocal microscope (Leica) at the Bioimaging Resource Center at Rockefeller University and quantified by mean fluorescence intensity relative to DAPI. Quantification was performed blind.

### Viral Production & Stable Cell Line Generation

Stable generation of cell lines was performed as previously described^1,6,7^. Lentivirus was generated using the ViraSafe lentiviral packaging system (Cell Biolabs) with Lipofectamine 2000 (Invitrogen) in HEK293T cells. Transductions were performed with 8 ug/mL polybrene. Plasmids to overexpress tRNA^Ile^ or with shRNAs targeting tRNA^Ile^_GAU_ were cloned into the plko.1 puromycin (Addgene # 8453) or blasticidin (Addgene #26655) backbone with AgeI/EcoRI restriction sites^22,23^. CRISPRi stable cells lines were generated with lentiviral transduction of dCas9-KRAB (Addgene # 110820) and pSLQ plasmid (Addgene # 51024) cloned with a tRNA^Ile^_GAU_ targeting guide (5’ TGAGCTAACCGGCCGCCCGA 3’), and then flow sorted for positive BFP+ and mCherry+ cells^24,25^. CRISPR generated cells were transduced with lentiCRISPRv2 (Addgene # 98290) cloned with specific guides targeting tRNA^Ile^_UAU_ loci (Guide 1: 5’ GCGCTAACCGATTGCGCCAC 3’, Guide 2: 5’ TGGCGCAATCGGTTAGCGCG 3’) or the eGFP targeting sequence as control (5’ GGGGCGAGGAGCTGTTCACCG 3’). Cells were then either selected with 2 ug/mL puromycin or 7.5 ug/mL Blasticidin (Thermo Fisher Scientific).

### Animal Studies

For metastasis assays, tail veins injections were performed in 5-6 week age matched female NOD SCID Gamma mice (The Jackson Laboratory #005557). Cells were counted via hemacytometer and resuspended in 1x PBS, and 100uL was injected with a 27G ½ needle (BD) into the lateral tail vein. Non-invasive bioluminescence imaging was performed immediately after injections using an IVIS Lumina II (Caliper Life Science) for Day 0 baseline, followed by weekly imaging. Bioluminesence imaging was obtained through retro-orbital injection of 50uL D-luciferin (Perkin Elmer) followed by 1 minute exposure in IVIS Lumina II. Unless otherwise stated, each experimental group consisted of n=5 mice. For bioluminescence imaging, cell lines were transduced with triple reporter and FACS sorted for GFP positive cells 48hours post transduction^4,10^. All animal work was conducted in accordance with protocols approved by the Institutional Animal Care and Use Committee at The Rockefeller University.

### Histology

Lungs were prepared by perfusion fixation with 4% paraformaldehyde through the circulation via the right ventricle post euthanasia. Lungs were then fixed in 4% paraformaldehyde overnight at 4°C. The samples were then embedded in paraffin and sectioned in 5 μm slices that were used for immunostaining. 5 μm sections at different depths were stained with hematoxylin and eosin (H&E).

### Ribosomal Profiling

Ribosomal profiling was performed based on the McGlincy & Ingolia protocol^4^. Briefly, cells were plated in 15 cm dishes at 50% confluency the day before collection. The media was aspirated, and the cells were washed with 5 mL ice cold PBS, and aspirated. The plate was then submerged in liquid nitrogen to freeze the cells. 400 uL ice cold lysis buffer was added to the plate, and scraped immediately. Each lysate was kept on ice until all plates were collected. Several plates were combined with lysis buffer totaling 1 mL per biological replicate. Lysates were then triturated ten times with a 26 gauge needle. Lysates were clarified at top speed for 10 mins in a cold bench top centrifuge and the supernatant was recovered and snap frozen in liquid nitrogen, then stored at −80 °C. Lysates were quantified with Quant-iT Ribogreen assay (Life Technologies) and 60 ug total RNA per sample was incubated with 3 uL RNase I (Epicentre #N6901K) for 45 minutes at RT with light shaking. 10 uL SUPERase*In RNase Inhibitor (Invitrogen) was added to stop digestion, and the RNA was transferred to a 13 mm x 51 mm polycarbonate ultracentrifuge tube. 900 uL Sucrose cushion (1 M Sucrose with 20 U/mL SUPERase*In in polysome buffer^4^) was underlaid and spun at 100,000 RPM at 4°C for 1 hour. With ribosomes pelleted, the supernatant was pipetted out of the tube. 300 uL Trizol was added to the pellet and resuspended. RNA was subsequently purified with the Direct-zol kit (Zymo). RNA was then precipitated overnight and resuspended after ethanol washes in 5 uL 10 mM Tris HCl pH 8.0. Ribosome footprints were isolated after running a 15% TBE-Urea gel and the RNA was excised within the range of 17nt – 34nt and then precipitated overnight. RNA fragments were then dephosphorylated with T4 PNK and ligated to a DNA linker with T4 Rnl2(tr) K227Q (NEB #M0351S) for 3 hours with distinct linker barcodes. Unligated linkers were depleted with yeast 5’-deadenylase (NEB #M0331S) and RecJ exonuclease (Epicentre #RJ411250) at 30°C for 45 minutes. Ligations were then purified with the Oligo Clean & Concentrater kit (Zymo) and samples were pooled. Ribo Zero Gold was then used to deplete ribosomal RNAs (2 reactions were used, and the 50°C step was omitted, Illumina). RNA was then purified using Oligo Clean & Concentration kit. The pooled ligations were then reverse transcribed using Superscript III at 55°C for 30 minutes, with RNA templates hydrolyzed by 2.2 uL 1 M NaOH. Samples were purified with the Oligo Clean & Concentrator kit and run on a polyacrylamide gel and the RT product was excised above 76nt. Gel slices were incubated with DNA gel extraction buffer overnight after the gel was broken up with gel breaker tubes (IST Engineering) and precipitated overnight. The RT product was resuspended in 10mM Tris HCl pH 8.0 and circularized with CircLigase II at 60°C for 1 hour. qPCR quantification of circulization products were performed to quantify number of cycles sufficient for library preparation, with the concentration estimated at 713 pM. 8 cycles were used to amplify the library with Pfusion with the primers indicated, NI-799 and NI-798^4^. Products were purified and size selected at >136bp, primarily at 160bp. The library was then precipitated, quality checked with Tapestation and sequenced on the NextSeq High Output 75 Single Read at the Rockefeller University Genomics Resource Center. Concurrently 1 ug total RNA was prepped for RNA sequencing according to the manufacturer’s instructions (Illumina). Analysis was performed as described previously^1^. For analysis, reads were first subjected to linker removal and quality trimming (cutadapt v1.14). The reads were then aligned against a reference database of rRNAs (iGenomes: AbundantSequenes) and tRNAs (GtRNAdb, hg38) so as to remove contaminants (using bowtie 2.3.4.1). STAR v2.5.2a was then used to align the remaining reads to the human transcriptome (build hg38). Xtail was used to count ribosome protected fragments, estimate translation efficiency, and perform statistical comparisons^26^.

### Polysome Profiling

Polysome profiling was adapted from Gandin et. al.’s protocol and with direction and assistance from Dr. Alison Ashbrook in Dr. Charlie Rice’s laboratory^27^. The day before cell collection, 7.5 million MDA-231 were plated in 15 cm plates (2 plates per experimental biological replicate) in normal DMEM media supplemented with 10% FBS. Cells were plated to achieve approximately 80% confluency at collection time to optimize polysome content. Each plate was treated for 5 minutes at 37 °C with DMEM with 100 ug/mL cycloheximide. The plate was then transferred to ice and the cycloheximide media was aspirated. Cells were washed twice with ice cold 1x PBS with 100ug/mL cycloheximide. All PBS was then aspirated carefully and the 15 cm plate was flash frozen in liquid nitrogen. 425 uL Lysis Buffer (5 mM Tris HCl pH 7.5, 2.5 mM MgCl2, 1.5 mM KCl, 100 ug/mL cycloheximide, 2 mM DTT, 0.5% Triton X-100, 0.5% sodium deoxycholate, and 100 units of SUPERase*In RNase Inhibitor (Invitrogen) 1x Protease Inhibitors EDTA-free) was then added to the plate and cells were scraped and transferred to an eppendorf tube on ice. Lysates were then spun at high speed at 4°C for 7 minutes to pellet nuclei. Supernatant was transferred to a new tube and the RNA concentration was measured using the Quant-iT Ribogreen assay (Life Technologies). 64ug RNA lysate was used for polysome fractionation. 10-50% Sucrose gradients were prepared the day before ultracentrifugation. Ultracentrifuge polyallomer tubes (Beckman Coulter, Cat #331372) were marked halfway and ~5.5 mL 10% sucrose polysome gradient buffer (20 mM Tris HCl pH 7.5, 140 mM KCl, 5 mM MgCl2, 10% Sucrose, 100 ug/mL cycloheximide, 0.5 mM DTT, 20 U/mL SUPERase*In) was added with a 10 mL sterile syringe to 1/8 inch above the line. 50% sucrose polysome gradient buffer (20 mM Tris HCl pH 7.5, 140 mM KCl, 5 mM MgCl2, 50% Sucrose, 100 ug/mL cycloheximide, 0.5 mM DTT, 20 U/mL SUPERase*In) was then underlain until the 10% sucrose layer was pushed above the marked line. The syringe was wiped with a Kimwipe prior to addition of 50% sucrose buffer to maintain separation between buffers. Black caps were added carefully to prevent the accumulation of bubbles in each ultracentrifuge tube. The Biocomp gradient master was then used at the following conditions: Long Cap 10% - 50% WV Step 1, 1:50 minutes, 80° angle, 21 speed. Gradients were then sealed with parafilm and incubated at 4°C overnight. Gradients were then balanced to within 10mg of each other. Normalized cell lysates were added (500 uL volume) and spun in a SW41 ultracentrifuge rotor at 38,000 RPM for 2 hours at 4°C. 60% sucrose was then used to fractionate spun lysates into 1 mL fractions and polysome peaks were measured with a Combi Flash UV-vis detector (Brandel) and TracerDAQ software. Polysome fractions were then pooled into appropriate groups: highly translated (Higher than 3 ribosomes, past the 1^st^ peak), and lowly translated (1-2 ribosomes and 80s) based on A280 UV peaks. The pooled fractions were then incubated with 3:1 Trizol LS Reagent, vortexed thoroughly, and incubated at RT for 5 minutes. RNA was then extracted following the instructions of the Direct-zol Miniprep Ki (Zymo Research), and eluted in 50 uL. RNA was quantified and normalized for input into either tRNACapture-seq qPCR with 5S, tRNA^Ile^_GAU_, or tRNA^Ile^_UAU_ probes (250 ng) or as Input into the QuantSeq 3’ mRNA-Seq Library Prep Kit (Lexogen) kit. 500 ng RNA was used as input and samples were processed according to QuantSeq (Lexogen) instructions. A pooled library was compiled using manufacturer’s primers and 10 nM Pool was quality checked with Tapestation and sequenced on the NextSeq High Output 75 Single Read at the Rockefeller University Genomics Resource Center.

For analysis, reads were mapped to the human transcriptome using STAR (v2.5.2a) with genome build hg38 and the number of reads for each gene was tabulated using featureCounts (v1.6.1). The Bioconductor package DESeq2 was then used to compare the fractions in control and tRNA^Ile^_GAU_/tRNA^Ile^_UAU_ modulated samples.

### Proteomics

Cells were lysed in 20 mM Tris HCl pH 8.0, 1% NP-40, 2 mM EDTA with 1x protease inhibitors (Roche). 50 ug lysate was used for label free quantification at the Rockefeller University Proteomics Core Facility. Maxquant software was utilized to compare three replicates per experimental group. Label free quantitation (LFQ) was used to compare the same peptide/protein between experimental groups (n=3 samples per group), which relies on normalization and strict filter criteria determined by the Proteomics Core. Student’s t-test difference and student’s t-test was then used to analyze the data.

### Immunofluorescence

Paraffin embedded histology slides from metastatic nodules were used. Slides underwent deparaffination via 5 minute incubations in xylene 2x, 100% Ethanol, 2x, 70% Ethanol, 50% Ethanol, and subsequently 1x PBS. Antigen retrieval was performed with 1x Citrate Buffer pH 6.0 in a microwave for 20 minutes. Slides were cooled to room temperature, then tissue regions were isolated with a PAP pen. Slides were then blocked with 10% Goat Serum (Sigma Aldrich) for 30 minutes at room temperature. Primary antibodies were incubated overnight at 4°C in a moist chamber (Vimentin V9 mouse (Abcam ab8069) 1:50), or for 2 hours at room temperature (Ki67 (Abcam ab927420) 1:200). Slides were washed with 0.5% Tween 20 PBS then incubated with secondary antibody for 1 hour at RT (1:200). Slides were then stained with DAPI for 5 minutes, and sealed with Prolong Gold anti-fade solution (Thermo Fisher). Fluorescent intensity was measured on an Inverted TCS SP8 laser scanning confocal microscope (Leica) at the Bioimaging Resource Center at Rockefeller University and quantified by number of positive cells per field of view. Quantification was performed blind.

### In Vitro Growth Assays

Cells at similar confluencies were resuspended in new DMEM media and counted with a hemacytometer. Cells were then seeded in equal numbers (100K for stress conditions, 50K for normal in vitro conditions) in 6 well plates in triplicate. Cells were then counted at the endpoint day with a hemacytometer. Each experiment was conducted three times. Cells treated with 200 uM H202 were counted on Day 3. Cells exposed to 0.5% hypoxia in an InvivO^2^ chamber (Baker Ruskinn) were quantified on Day 3. Growth assays in normal *in vitro* conditions were quantified on Day 5.

### Western Blot

Cells seeded a day previously were washed with 1x PBS and then lysed with either RIPA buffer or 20 mM Tris HCl pH 8.0, 1% NP-40, 2 mM EDTA with 1x protease inhibitors (Roche). Protein concentrations were quantified with a BCA Kit (Thermo Fisher) and normalized. Protein lysates were run at 200V for an hour through either a 4-12% Bis-Tris or 3-8% Tris Acetate gel (Invitrogen), and then transferred at 300 mA for one hour in 15% methanol 1x Transfer Buffer on a methanol activated PVDF membrane. Membranes were then stained with Ponceau and blocked for one hour in Odyssey^®^ Blocking Buffer. Primary antibody incubations occurred overnight at 4°C on a rocker at the following concentrations: alpha tubulin 1:1000, (Proteintech) SMNDC1 1:500 (Proteintech). Membranes were then washed with 0.05% Tween 20 PBS three time and incubated with mouse or rabbit fluorescent IRDye^®^ conjugated secondary antibodies 1:20,000 (LI-COR Biosciences) for one hour. Membranes were subsequently washed three times and imaged and quantified using the Odyssey^®^ Sa Infrared Imaging System at the Rockefeller University Center for High Throughput Screening. Quantification was done using the Image Studio Lite^TM^ software.

### Codon Reporters

Wildtype or codon mutant SMNDC1 (all AUA codons changed to AUC codons) coding sequence gene blocks were designed with NheI & XhoI restriction sites and a N-terminal flag tag and ordered from IDT. SMDNC1 gene blocks were cloned into the psiCheck 2 vector. The firefly luciferase was removed and replaced with a renilla luciferase with only AUU encoding isoleucines via restriction cutting with PspOMI & XbaI. This adapted SMNDC1 reporter was transfected with 2.5 ug plasmid and 10 uL Lipofectamine 2000 (Thermo Fisher) in triplicate in MDA-MB-231 cells with modulated tRNA^Ile^ levels. Cells were lysed after 24 hours and protein was extracted with RIPA buffer with 1x protease inhibitors EDTA-free (Roche). Protein expression was measured through LICOR Western blotting as described above.

### RSCU and Pathway Enrichment Analyses

#### Gene filtering

The Homo sapiens GRCh38 CDS sequences were downloaded from the Ensembl database. To avoid multiple splice variants from the same gene affecting downstream analysis, principle splice isoforms were filtered using annotations from the APPRIS database and a custom Python script. For genes with multiple annotated isoforms, the transcript with the highest score was chosen as the representative.

#### RSCU calculation

To calculate the relative synonymous codon usage (RSCU) for a given codon in each gene, we first calculated the total abundances of each codon across our entire filtered CDS dataset to determine the empiric distribution of synonymous codon usage for each amino acid. For each gene, the RSCU score was calculated as: [Observed_Codon_Usage – Expected_Codon_Usage] / Expected_codon_usage. The expected codon usage was defined as Observed_Amino_Acid_Usage * Pr(Codon_Usage | Amino_Acid) where the probability mass function was determined using the empiric codon distribution described above. For genes/transcripts in which a given amino acid appeared zero times, the RSCU score was set to 0.

#### Mutual Information/Pathway Enrichment

Genes were ranked by the RSCU score calculated above. Mutual information analyses to detect significantly over-represented and under-represented pathways in discrete bins were performed using the iPAGE mutual information framework with pathway annotations built from the Reactome database^28^. Heatmaps were generated with iPAGE. To determine pathways that were most likely to be divergently modulated by AUA or AUC over/under-expression, we additionally filtered the output to include pathway enrichments/depletions present in both AUC and AUA analyses with p-values less than 10E-3 in the highest RSCU bin. Heatmaps were generated using Python software and the seaborn package.

##### Databases/Sites/Software used

Ensembl: https://useast.ensembl.org/index.html

APPRIS: http://appris-tools.org/#/

iPAGE: https://tavazoielab.c2b2.columbia.edu/iPAGE/

Reactome: https://reactome.org

Python 3.6.0: https://www.python.org

Pandas: https://pandas.pydata.org

Seaborn: https://seaborn.pydata.org

#### Statistical analysis

Results are presented in dot-plot with dots representing individual values and bar-charts depicting average values with standard error of the mean (±s.e.m.). The number of samples for each group was chosen based on the expected levels of variation and consistency. FISH quantification was performed in a blinded fashion. Unless otherwise stated, statistical significance was assessed by a two-tailed Student’s t-test with *P*-value < 0.05 being considered statistically significant.

## Data availability

Experimental data will be available from the corresponding author upon request. Sequencing data will be made available in public databases.

## Ethical regulations

All animal experiments were performed under supervision and approval of the Institutional Animal Care and Use Committee (IACUC) at the Rockefeller University.

## Acknowledgements

We thank members of the Tavazoie laboratory and Hani Zaher for thoughtful comments on previous versions of the manuscript. We thank Alison Ashbrook and Charlie Rice’s laboratory for technical assistance with polysome profiling. We also thank Rockefeller University resource centers: Alison North and staff at the Bio-Imaging resource facility, Connie Zhao from the genomics resource center, Soren Heissel and Henrik Molina from the proteomics resource center, and Vaughn Francis from the Comparative bioscience center and veterinary staff for animal husbandry and care. L.N and M.C.P. were supported by a Medical Scientist Training Program grant from the National Institute of General Medical Sciences of the National Institutes of Health under award number T32GM007739 to the Weill Cornell/Rockefeller/Sloan Kettering Tri-Institutional MD-PhD program. H.G. was supported by R00CA194077 and R01CA24098. S.F.T. is an HHMI Faculty Scholar, and was supported by the Breast Cancer Research Foundation award, the Reem-Kayden award, and by NCI grant R01CA215491. S.F.T. and the Tavazoie lab were supported by the Black Family and the Black Family Metastasis Research Center.

## Author Contributions

L.N., H.G. and S.F.T. designed the experiments. L.N, N.M., and M.C.P. performed the experiments. H.G. and H.A. performed ChIP-Seq, TGIRT, Ribosomal Profiling, Polysome Profiling sequencing, and ribosomal dwelling time analyses. D.H. performed iPAGE codon analyses. L.N. and S.F.T. wrote the paper with input from the co-authors.

---

**Supplementary Figure 1.**
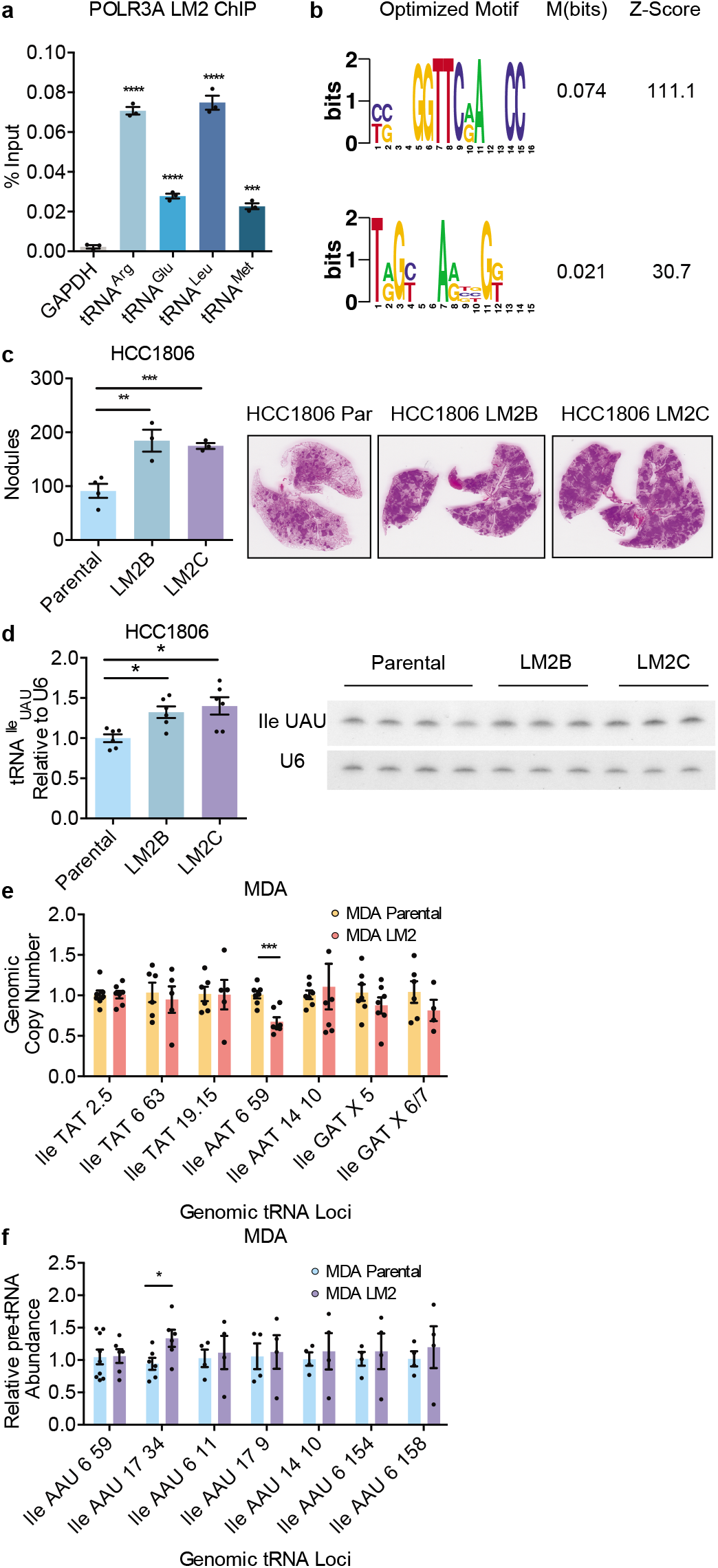
POLR3A ChIP sequencing reveals differential occupancy in isogenic poorly and highly metastatic breast cancer pairs. (a) tRNA genomic loci abundance measured as percent (%) input using RT-qPCR of POLR3A IP cDNA with GAPDH as a negative control. (b) Motif analysis of POLR3A IP normalized to Input sequences using FIRE analysis. (c) Quantification of lung metastatic nodules post extraction after tail vein injection of 1.5×10^5^ HCC1806 Parental or highly metastatic derivatives LM2B or LM2C, with representative histology; n=3-4 in each cohort. (d) Northern blot quantification of tRNA^Ile^_UAU_ relative to U6 of two independently derived LM2 lines relative to HCC1806 Parental cells with representative blot. (e) Relative genomic copy number of tRNA^Ile^ loci, quantified by RT-qPCR. (f) Relative pre-tRNA abundance of tRNA^Ile^_AAU_ across multiple primers covering distinct genetic loci using RT-qPCR of MDA LM2 vs. MDA-MB-231. Two-sided unpaired student’s t-tests performed, p-values represented *, **, ***, **** as p<0.05, p<0.01, p<0.001, p0.0001, respectively.

**Supplementary Figure 2.**
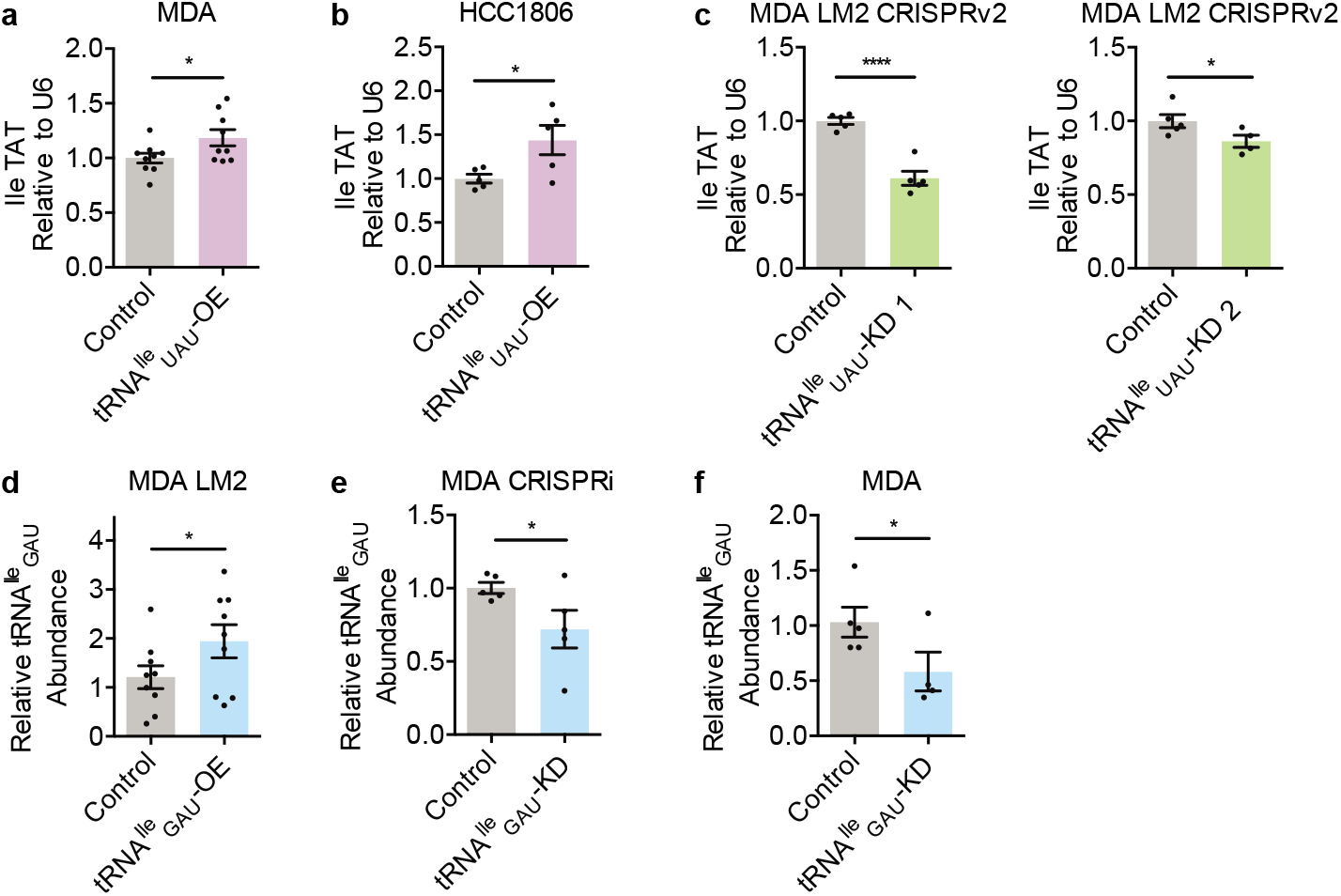
tRNA^Ile^_UAU_ and tRNA^Ile^_GAU_ levels can be manipulated exogenously. (a,b) Northern blot quantification of tRNA^Ile^_UAU_ relative to U6 of MDA (a) or HCC1806 (b) Parental cells with control or overexpression of tRNA^Ile^_UAU_. (c) Northern blot quantification of tRNA^Ile^_UAU_ relative to U6 of LM2 cells depleted of tRNA^Ile^_UAU_ via CRISPR with Guide 1 or 2 versus control. (d) tRNA^Ile^_GAU_ quantification by specific tRNA^Ile^_GAU_ probe RT-qPCR normalized to 18S probes of LM2 cells with control or overexpression of tRNA^Ile^_GAU_. (e,f) tRNA^Ile^_GAU_ quantification by specific tRNA^Ile^_GAU_ probe RT-qPCR normalized to 18S probes MDA Parental CRISPRi cells with guides targeting either control or tRNA^Ile^_GAU_ (e) or or shRNA targeting control or tRNA^Ile^_GAU_ (f). Two sided un-paired student t-tests performed, p-values representated as *, **, *** as p<0.05, p<0.01, p<0.001.

**Supplementary Figure 3.**
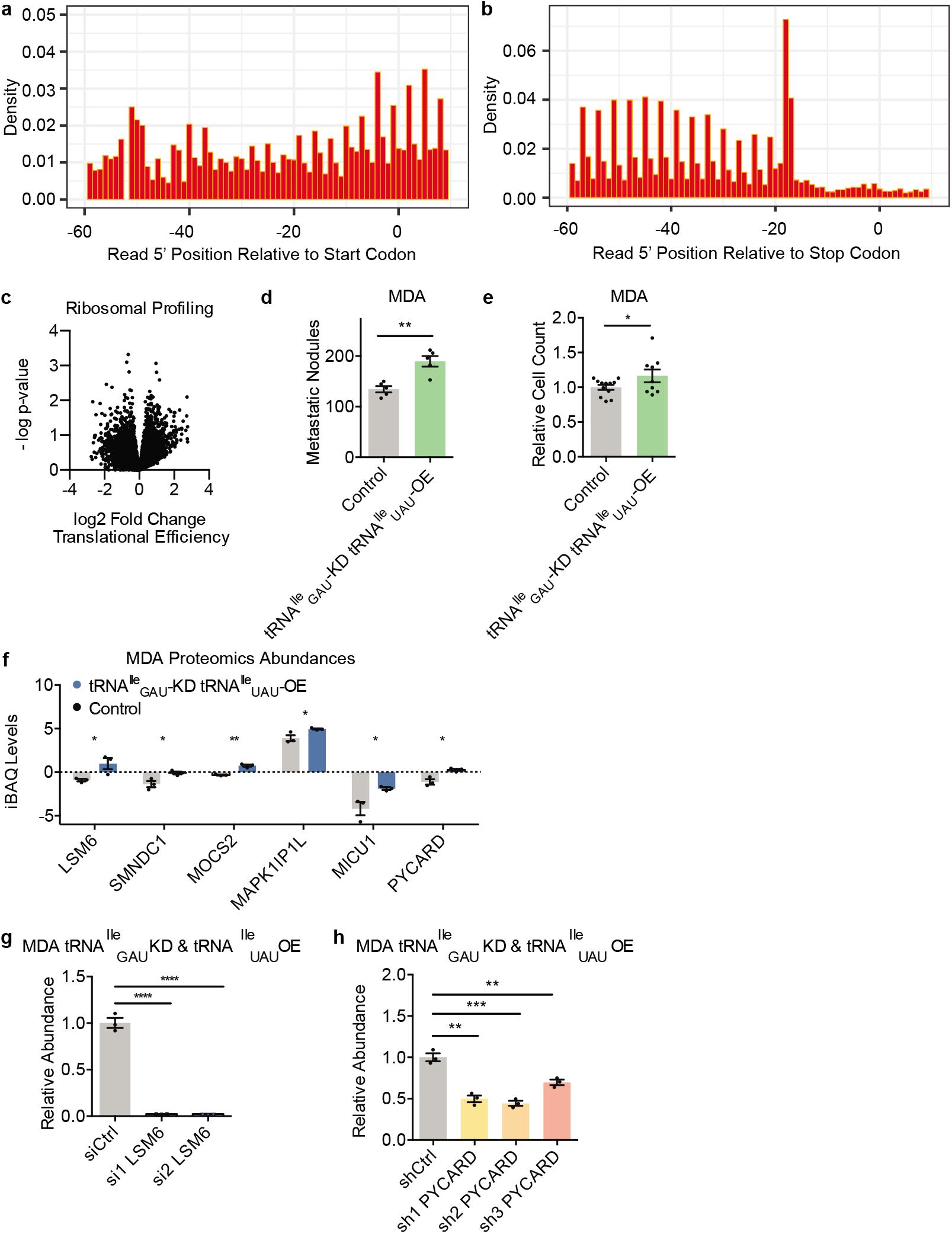
Downstream effectors of tRNA^Ile^_GAU_ depletion and tRNA^Ile^_UAU_ overexpression mediate increased growth under metastatic stress conditions. (a, b). Histogram of read coverage to demonstrate 3 nucleotide periodicity of the coding sequence with respect to the start (a) and stop codon (b) of the reading frame. (c) Volcano plot representing log2 fold change vs. –log p value of translational efficiency from ribosomal profiling of MDA tRNA^Ile^_GAU_ depleted and tRNA^Ile^_UAU_ overexpression cells versus control. (d) Quantification of lung metastatic nodules post extraction after tail vein injection of 1×10^5^ MDA tRNA^Ile^_GAU_ depleted and tRNA^Ile^_UAU_ overexpression cells versus control. (e) Relative cell counts of MDA MB 231 control & tRNA^Ile^_GAU_ depletion tRNA^Ile^_UAU_ overexpression cells after 5 days. Two-sided un-paired student’s t-tests performed, p-values represented as *, ** as p<0.05, p<0.01, respectively. (f) iBAQ values of six candidate downstream effectors, measured by label free quantification mass spectrometry; 3 biological replicates each. (g) RT-qPCR quantification of LSM6 cDNA levels normalized to GAPDH on Day 2 of siRNA transfection. (h) RT-qPCR quantification of PYCARD cDNA levels normalized to GAPDH. Two-sided un-paired student’s t-tests performed, p-values represented **, ***, **** as p<0.01, p<0.001, p,0.0001, respectively.

**Supplementary Figure 4.**
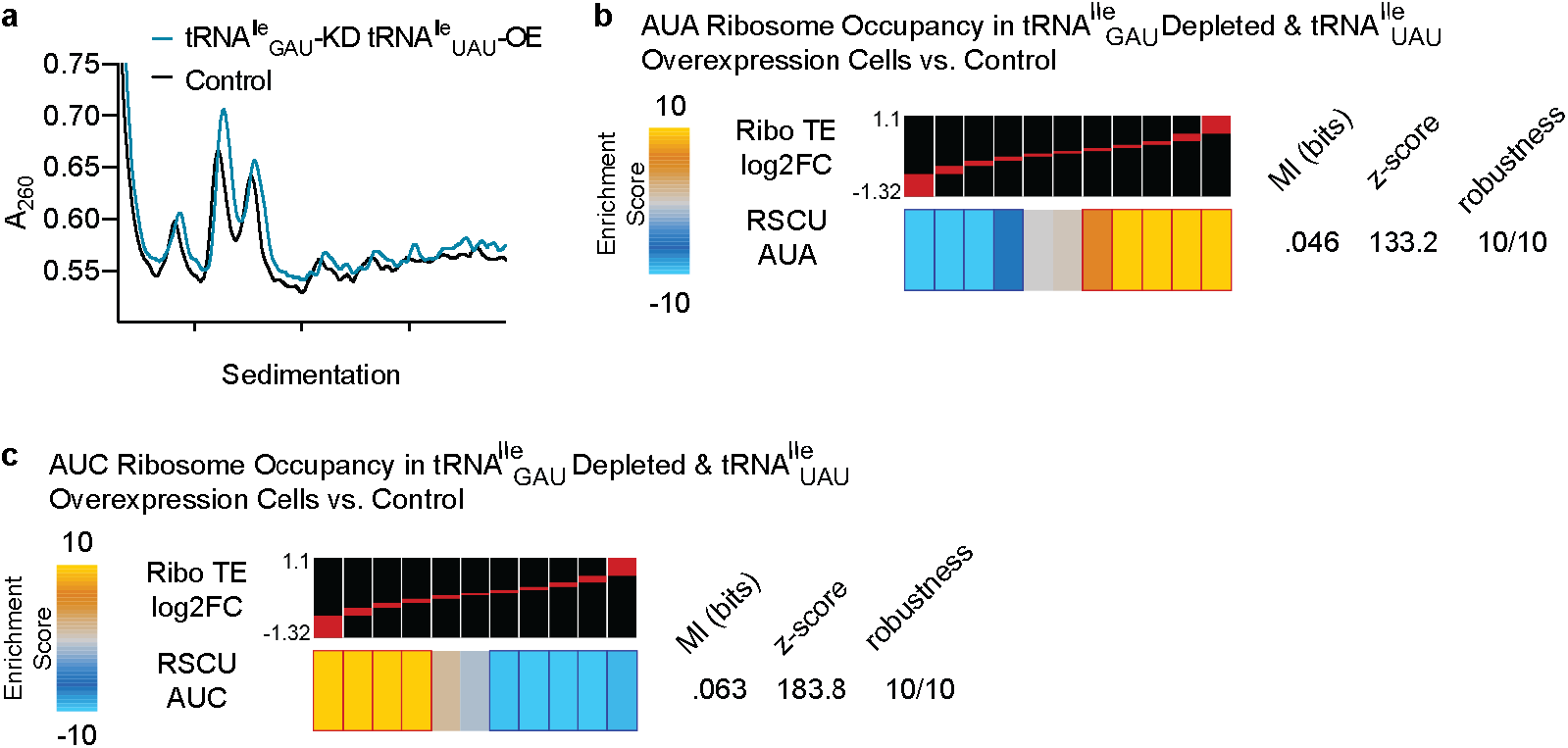
Ribosomal profiling of MDA cells concurrently modulated with tRNA^Ile^_GAU_ depletion & tRNA^Ile^_UAU_ overexpression. (a) Polysome traces of two representative samples, measured by UV spectrometry. (b) Genes with a high abundance of AUA codons were significantly enriched among genes significantly upregulated in ribosomal protected fragments (corrected for their transcript changes) in tRNA^Ile^_GAU_ depleted tRNA^Ile^_UAU_ overexpression cells versus control MDA MB 231 cells, measured by ribosomal profiling. The statistical significance of these enrichments was assessed using mutual-information calculations and associated Z score (based on randomized input vectors). Also included is the χ^2^ p value for the associated contingency table. The heatmap was generated using the –log of the hypergeometric p value for enrichment and log of p value for depletion (collectively termed the enrichment score). The red and dark-blue borders indicate the statistical significance of the calculated hypergeometric p values. (c) Same as (b) except analyzed for AUC codon enrichment. Codon content scored for by relative synonymous codon usage score (RSCU).

